# The β-Carotene-Oxygen Copolymer: its Relationship to Apocarotenoids and β-Carotene Function

**DOI:** 10.1101/2020.12.29.424736

**Authors:** Trevor J. Mogg, Graham W. Burton

## Abstract

β-Carotene spontaneously copolymerizes with molecular oxygen to form a β-carotene-oxygen copolymer compound (“copolymer”) as the main product, together with small amounts of many apocarotenoids. Both the addition and scission products are interpreted as being formed during progression through successive free radical β-carotene-oxygen adduct intermediates. The product mixture from full oxidation of β-carotene, lacking both vitamin A and β-carotene, has immunological activities, some of which derive from the copolymer. However, the copolymer’s chemical makeup is unknown. A chemical breakdown study shows the compound to be moderately stable but nevertheless the latent source of many small apocarotenoids. Although the copolymer alone is only slightly affected by heating at 100°C for 4 h, in methanol solution it is significantly degraded by hydrochloric acid or sodium hydroxide, liberating many apocarotenoids. GC-MS analysis with mass-spectral library matching identified a minimum of 45 structures, while more than 90 others remain unassigned. Thirteen products are Generally Recognized as Safe (GRAS) human flavor agents. Newly identified products include various small keto carboxylic acids and dicarboxylic acids, several of which are central metabolic intermediates. Also present are the dialdehydes glyoxal and methyl glyoxal, recently reported as β-carotene metabolites in plants. Although both compounds at higher concentrations are known to be toxic, at low concentration methyl glyoxal has been reported to be potentially capable of activating an immune response against microbial infection. In plants, advantage is taken of the electrophilic reactivity of specific apocarotenoids derived from β-carotene oxidation to activate protective defenses. Given the copolymer occurs naturally and is a major product of non-enzymatic β-carotene oxidation in stored plants, by partially sequestering apocarotenoid metabolites the copolymer may serve to limit potential toxicity and maintain low cellular apocarotenoid concentrations for signaling purposes. In animals the copolymer may serve as a systemic source of apocarotenoids.

## Introduction

The propensity of oxygen to preferentially add polymerically to highly unsaturated hydrocarbon compounds was demonstrated in the pioneering styrene oxidation model study published by Miller and Mayo in 1956.^1^ Yet, despite the presence of the highly unsaturated conjugated polyene backbone and many studies carried out over several decades,^2–4^ only relatively recently has it been reported that the spontaneous oxidation of β-carotene is dominated by the formation of a β-carotene-oxygen copolymer product (“copolymer”).^5^ Previously, the oxidation reaction had been characterized by the formation of a mixture of mostly well-known β-apocarotenoid products, which apparently arose from individual cleavages of the double bonds in the backbone. The copolymer product apparently escaped notice.

Full oxidation of β-carotene with air or pure oxygen in ethyl acetate solvent gives a reproducibly complex product, OxBC, containing 80-85% by weight of copolymer compound and 15-20% of many small apocarotenoid cleavage compounds.^5^ Here, the term apocarotenoid is applied somewhat loosely to also include additional small molecule compounds, the result of further chemical transformations. These carotenoid-derived compounds are also referred to in the literature less commonly and more generally as norisoprenoids.^6^

In OxBC the apocarotenoids contain no more than 18 carbon atoms, less than half of β-carotene’s 40 carbons.^5^ Notably, apocarotenoids formed early in the oxidation and containing more than 18 carbons, including vitamin A, are absent in OxBC, having undergone further oxidation. The several most abundant apocarotenoids are present at levels of ~1-2% or less by weight in OxBC made by air-oxidation, being slightly lower than for OxBC made with pure oxygen.^5^

Many of the smaller apocarotenoids are well known for their flavor and fragrance characteristics,^6^ being present, for example, in leaf products (e.g., tobacco, tea, mate), many essential oils, fruits (grapes, passionfruit, starfruit, quince, apple, nectarine), vegetables (tomato, melon), spices (saffron, red pepper), wine, rum, coffee, oak wood, honey and seaweeds. Several of the apocarotenoids present in OxBC have been designated as Generally Recognized As Safe (GRAS) human flavor agents.^7^

In plants there has been much recent research activity on the oxidative metabolism of β-carotene and the involvement of apocarotenoids in photosynthetic tissues^8–14^ and non-green Arabidopsis roots.^15–16^ Several β-apocarotenoids, e.g., β-cyclocitral, β-ionone, dihydroactinidiolide and β-cyclogeranic acid, have been identified as stress signals, which at very low concentrations can activate processes that protect plants.^12, 17^

Substantial β-carotene losses occur by non-enzymatic oxidation during storage of crops, fruits and vegetables, e.g. maize grains,^18^ and provitamin A-biofortified Golden Rice.^19–20^ Perhaps not surprisingly, given its spontaneous formation from β-carotene in air, the copolymer occurs naturally as a degradation product in a variety of plant tissues upon drying^21^ or storage.^20^ During rice endosperm development it has been reported β-carotene losses yield only low amounts of β-apocarotenoids, while the majority of β-carotene autoxidizes to form copolymers.^20^

In livestock animals, feeds supplemented with OxBC at parts-per-million levels provide benefits in poultry,^22^ swine^23^ and dairy cattle.^24^ The results are consistent with an immunomodulatory action.^25–26^ Because OxBC contains no vitamin A or β-carotene^5^ and shows no vitamin A activity,^25^ the activity is independent of vitamin A or of any effect of β-carotene alone. *In vitro* evidence points to the copolymer as a source of innate immune activity,^25^ whereas it is not yet known if the copolymer itself or the apocarotenoid fraction is responsible for an ability to modulate inflammatory response.^26^

To gain a better understanding of the copolymer in relation to β-carotene function in animals and plants, it is important to know more about the copolymer’s chemical properties and makeup, beyond the fragmentary information presently available.^5, 21^ Such knowledge bears on the question of the mechanism of the copolymer’s action upon immune function and its metabolism in animals. The copolymer has proved refractory to application of state-of-the-art NMR and mass spectrometry (MS) instrumentation, including 1D and 2D NMR and MALDI mass spectrometry techniques. The NMR results were limited to indicating a complex polymeric structure containing HC=CH, -CH=O, >CH-OH, -CH_2_-OH, >CH_2_, and -CH_3_ groups.

In this study, chemical breakdown studies on the copolymer compound have been carried out to assess its stability, to learn more about its chemical makeup and to identify possible metabolic breakdown products. Several conditions were employed, including the effect of solvents alone, acids, bases, oxidizing agents and heat, to determine the susceptibility of the copolymer to decomposition into its constituent components. The reaction products were evaluated by GPC, HPLC and GC-MS to identify changes to the copolymer itself and to identify, as much as possible, the small molecule compounds released.

## Materials and Methods

### Materials

The solid β-carotene-oxygen copolymer compound was obtained by successive solvent precipitations from OxBC, as has been described previously.^5, 21^ Solvents of analytical or HPLC grade were used in all experiments.

### General

GPC-UV chromatograms were obtained using an HP 1090 HPLC apparatus equipped with a diode array detector and a 7.8 × 300 mm Jordi Flash Gel 500A GPC column (5 μm particle size; Jordi Laboratories LLC, Bellingham, MA 02019 USA). Samples were dissolved in and eluted with tetrahydrofuran at 1 mL/min for 14 min.

HPLC-UV chromatography was carried out using an Agilent HP Series II 1090 with UV diode array detector. Analysis of the oxidation product mixtures was carried out with a Waters C18 Atlantis T3 column (4.6 mm x 100 mm, 3 μm) with a guard column (C18 Atlantis T3, 4.6 mm x 20 mm, 3 μm). Samples were dissolved in acetonitrile. The following conditions were used: solvent A, water; solvent B, acetonitrile; flow 1 mL/min; column equilibrated with 95/5 A/B mobile phase for 5 minutes; gradient elution: 5%-100% B over 0-15 min then held for 10 min.

GC–MS was performed with an Agilent Technologies 6890N GC with a 5975B VL mass selective detector operated in electron ionization mode. The GC was equipped with an HP 5 column, 30 m × 0.25 mm × 0.25 μm. Measurement conditions: initial pressure 17 psi, constant flow of 1.0 mL/min; injector temperature 250 °C; initial oven temperature 50 °C for 1 min, temperature ramp 20 °C/min to 280 °C, hold time 2.5 min. For compound identification, spectra were acquired in scan mode and compared to the NIST 05 Spectral Library.

### Copolymer breakdown studies

The experiments are listed in Table 1. Room temperature experiments were carried out in sealed 20 mL glass scintillation vials without stirring. All other experiments were carried out with stirring in round bottom flasks and were open to the atmosphere. In experiments conducted at reflux temperatures, a condenser was attached to the flask, which was heated by placing in an oil bath. Concentrated hydrochloric acid (12.1 M), glacial acetic acid, sodium hydroxide pellets, and 30% aqueous hydrogen peroxide were mixed directly into methanol, then added to the copolymer.

**Table 1.**
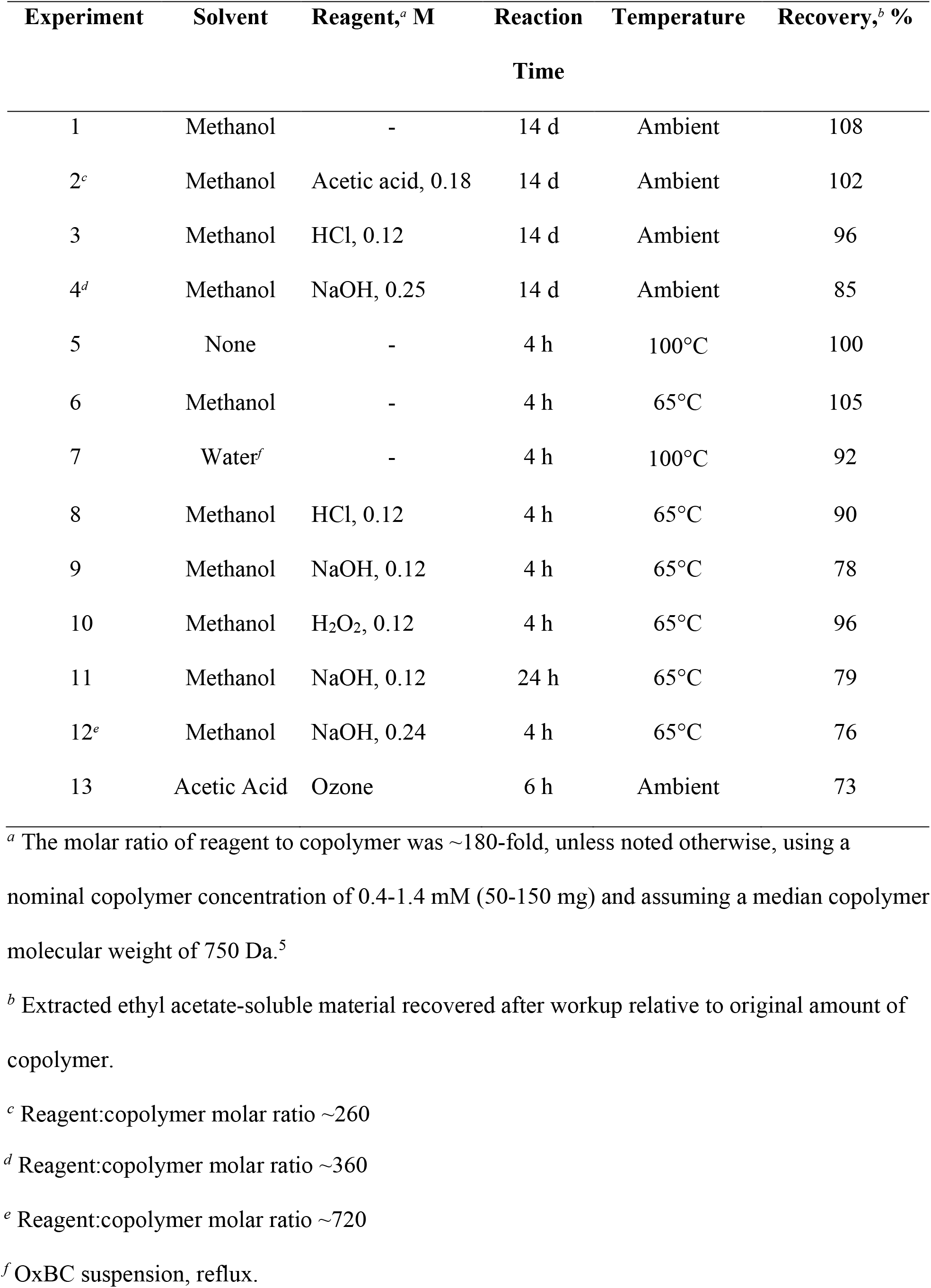
Summary of chemical and thermal degradation experiments of the OxBC copolymer and associated product recoveries.

#### Workup procedures

Methanol solvent alone (Expts. 1, 6): the solvent was removed by rotary evaporation, the residue taken up into ethyl acetate, the solution rotary evaporated, and the residue dried under vacuum. Copolymer alone without solvent (Expt. 5): the sample was taken up in ethyl acetate, rotary evaporated and the residue dried under vacuum. Hydrochloric acid or acetic acid/methanol (Expts. 2, 3, 8): the solvent was rotary evaporated, brine was added (10 mL) to the residue and the mixture extracted with ethyl acetate (4 x 10 mL). The combined organic extracts were washed once with brine (10 mL), dried over sodium sulfate, filtered, rotary evaporated, and the residue dried under vacuum. Sodium hydroxide/methanol (Expts. 4, 9, 11, 12): the procedure was the same as for the acidic samples, except aqueous HCl (1 M) was first added to acidify the solution to pH ~ 2. Water suspension reflux (Expt. 7): the sample heated in water was saturated with sodium chloride, extracted with ethyl acetate (4 x 10 mL), and the combined extracts dried over sodium sulfate, filtered, rotary evaporated and the residue dried under vacuum. Hydrogen peroxide/methanol (Expt.10): the methanol was evaporated, brine (10 mL) added, and the mixture extracted with ethyl acetate (4 x 10 mL). The combined extracts were washed with brine (10 mL), dried over sodium sulfate, filtered, rotary evaporated and the residue dried under vacuum.

#### Ozonolysis

A sample of copolymer (153 mg) dissolved in glacial acetic acid (10 mL) was ozonized at room temperature using pure oxygen gas and an ozone generator (Type: 1VTT, Ozomax Inc., Granby, Quebec). After ozone was bubbled through the solution for 6 hours, nitrogen gas was bubbled through the solution for 10 min, then hydrogen peroxide (30% aq., 7 mL) was added and stirred overnight. The reaction was heated to reflux for 1.5 hours and then water and acetic acid were removed by rotary evaporation. The residual colourless oil was dissolved in ethyl acetate (6 mL), dried over sodium sulfate, filtered, and the solvent evaporated. The product was dried for 20 min under vacuum to give a viscous, colourless oil (73% yield). A sample of the recovered product was methylated with trimethyloxonium tetrafluoroborate and analyzed by GC-MS. A blank ozonolysis without copolymer was carried out and analyzed to exclude any compounds not originating from the copolymer (e.g., from gas tubing).

A sample of lycopodium sporopollenin (50 mg in 10 mL acetic acid; Polysciences Inc., Warrington PA) was ozonized in a similar manner, except that ozone was bubbled for 10 hrs. At the start, the reaction mixture was a dark brown suspension, and at the end it was pale yellow with a small amount of precipitate. Hydrogen peroxide (30% aq., 2 mL) and water (2 mL) were added and stirred overnight. By morning the precipitate had disappeared, leaving a clear, pale yellow liquid. More hydrogen peroxide (30% aq., 1 mL) was added, and the mixture refluxed 1.5 hrs. The mixture was rotary evaporated to give a yellow oil (65 mg). A sample of the oil was methylated with trimethyloxonium tetrafluoroborate and analyzed by GC-MS.

#### Methylation

For esterification of acids into methyl esters, a sample of reaction product (ca. 10 mg) was dissolved in MeOH (4.5 mL) and aq. NaHCO_3_ (1 M, 1 mL) was added. The mixture was stirred and trimethyloxonium tetrafluoroborate (95%, Sigma-Aldrich, Oakville, ON) was added in 3 portions over 5 min for a total of ca. 0.3 g. The mixture was stirred for 10 min. and maintained weakly basic by ensuring the presence of solid NaHCO_3_, adding small amounts as needed. Water (6 mL) was then added, and the mixture extracted with dichloromethane (2 x 6 mL). The combined extracts were dried over sodium sulfate, filtered, and rotary evaporated. The residue was dissolved in acetonitrile for GC-MS analysis.

### Determination of geronic acid (GA) content of copolymer breakdown product mixtures

The basic GA analysis procedure used for the copolymer and its breakdown product mixtures has been described previously.^5, 21^ Samples were spiked with deuterium-labelled GA (GA-d_6_), purified by passage through a solid-phase extraction cartridge, and esterified with trimethyloxonium tetrafluoroborate prior to GC-MS analysis.

Copolymer or breakdown product mixtures (10-25 mg) were placed in a 20 mL vial with stir bar and dissolved in MeOH (3 mL). GA-d_6_ solution (0.4 mL, 0.1694 mg/mL in MeOH) was added. The mixture was stirred for 1 min and the solvent evaporated under a stream of N_2_. Aqueous NH_3_ (5%, 6 mL) and water (3 mL) were added, the vial capped, and the solution stirred vigorously for 20 min. SPE cartridges (Waters Oasis MAX, 500 mg/6 mL) were prepared by the passage, in sequence, of methanol (6 mL), water (6 mL) and aqueous NH_3_ (0.5%, 4.5 mL). The basic analyte solution was passed through the cartridge under gravity, the cartridge washed with aqueous NH_3_ (0.5%, 4.5 mL) followed by methanol (9 mL). The carboxylic acids were eluted by passage of a solution of HCl in MeOH (2%, 4.5 mL) and collected in a 20 mL vial. Solid NaHCO_3_ was added and stirred until bubbling ceased, followed by addition of NaHCO_3_ (1 M, 1 mL) to give a cloudy solution. The solutions were stirred gently while Me3OBF_4_ was added in three portions (total ~ 0.2 g), then stirred vigorously for 15 min and maintained slightly basic by adding a small amount of solid NaHCO_3_. After 15 min, H_2_O (6 mL) was added, stirred, and the mixture extracted with CH_2_Cl_2_ (2×7 mL). The combined CH_2_Cl_2_ extracts were dried (Na_2_SO_4_), transferred by pipette to a small flask and solvent carefully evaporated at room temp to minimize loss of volatile methyl geronate. The residue was dissolved in acetonitrile (0.6 mL), passed through a 0.2 μm syringe filter and 1 μL injected into the GC-MS instrument. GC-MS calibration and analysis of GA was carried out as described before.^5, 21^

## Results

The stability of the copolymer and the nature of its chemical makeup were addressed in a series of experiments that subjected the compound to individual treatments with heat and an excess of acidic, basic and oxidizing agents. Methanol was used as solvent because the copolymer is insoluble in water. Experiments were conducted at room temperature for 14 d and at reflux (65°C) for 4 h or 24 h. The polymer itself was also heated without solvent at 100°C for 4 h. The product recoveries listed in Table 1, obtained after sample workup, represent the sum total of lipid-soluble material plus any remaining intact copolymer that was recoverable by extraction into ethyl acetate, and do not include any water-soluble products retained in the aqueous phase. The material recovered in the ethyl acetate extracted fraction of the reaction product mixture from each experiment was evaluated by GPC and HPLC to identify any changes in the copolymer and by GC-MS to identify, where possible, any degradation products that were formed.

### Copolymer stability

As a starting point, samples of OxBC copolymer were dissolved in methanol with or without acid or base and stood at room temperature for 14 d (Table 1, Expts. 1-4).

GPC results are shown in Fig. 1. The untreated, intact copolymer begins to elute at ~5.6 min and displays a symmetric peak centered at ~7.7 min (Fig. 1A). The effect of exposure to solvent and reagents caused varying amounts of partial copolymer decomposition, which manifested as an increase in GPC elution time because of a decrease in mean molecular weight. There was an accompanying loss of peak symmetry and an appearance of several small low molecular weight peaks on the trailing edge of the copolymer peak, corresponding to the release of small molecule compounds.

**Fig. 1.**
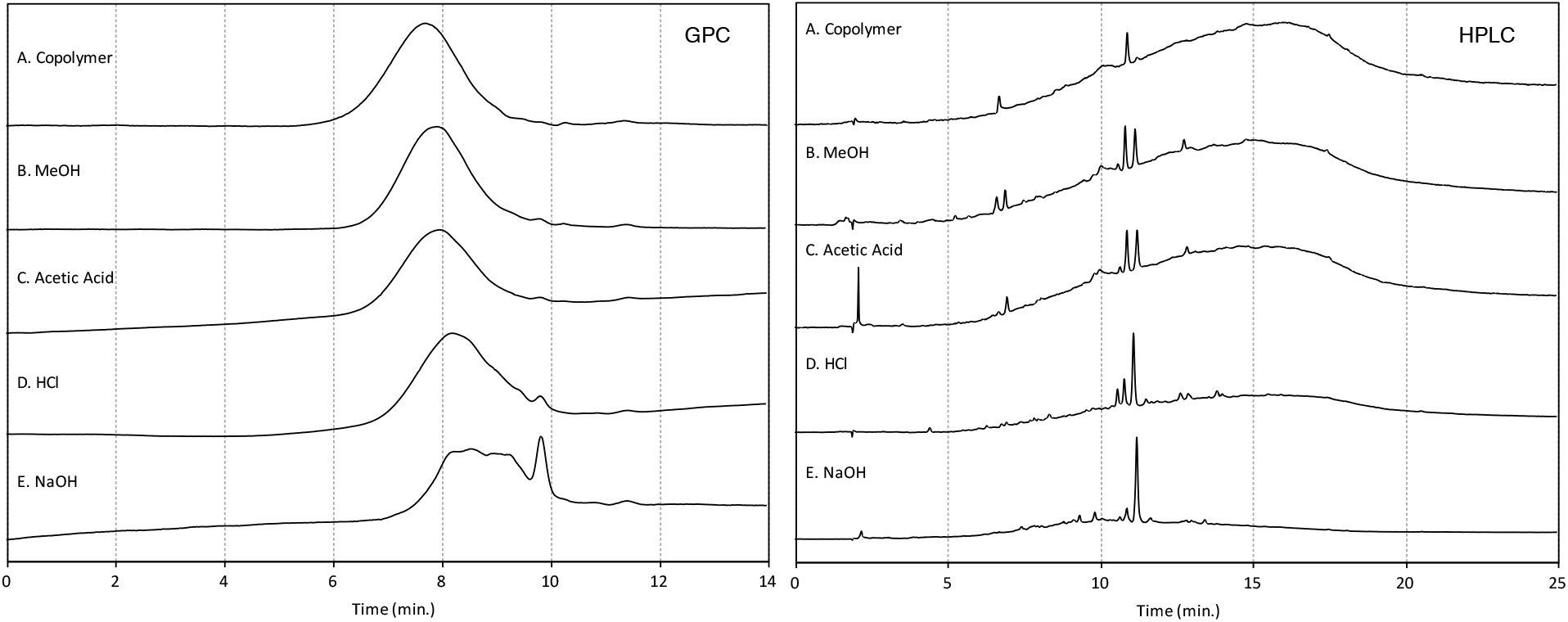
GPC-UV and HPLC-UV chromatograms at 220 nm of the copolymer reference compound (A) and of ethyl acetate extracts of products obtained from methanol solution of copolymer after standing for 14 d at room temperature in methanol alone (B) or treatment with acetic acid (C), hydrochloric acid (D) and sodium hydroxide (E), respectively. (Expts. 1-4, Table 1). Injected sample sizes were ~200 μg (GPC A, B, D; HPLC A-E) and ~100 μg (GPC C, E).

Incubating the copolymer compound in methanol solution or in an excess of acetic acid in methanol for 14 d at room temperature both resulted in the appearance of minor amounts of cleavage products and a small reduction in the copolymer’s mean molecular weight, with the peak eluting at ~7.9 min (Figs. 1B, 1C). An excess of hydrochloric acid promoted greater breakdown, with a less symmetrical polymer peak, displaced to ~8.2 min, together with a small breakdown product peak clearly evident at ~9.8 min (Fig. 1D). An excess of sodium hydroxide caused the largest change. Elution of the copolymer was further delayed, and the peak profile was irregular, with the main peak flattened and a prominent breakdown product peak eluting at ~9.8 min (Fig. 1E).

The HPLC chromatograms corroborated the GPC findings. The chromatogram of the intact copolymer by itself is dominated by a large, asymmetric rise in the baseline as the copolymer elutes, with several small peaks of cleavage products superimposed on the leading edge (e.g., at 6.7 and 10.8 min) (Fig. 1A). It is likely that during purification the copolymer was not entirely freed of contaminating cleavage compounds originally present in OxBC. Several additional breakdown product peaks appeared after standing for 2 weeks in methanol or with acetic acid in methanol, e.g., at 6.9 and 11.2 min. (Figs. 1B, 1C), although there was no discernible change in the extent and magnitude of the baseline rise compared to the copolymer (Fig. 1A). However, it is apparent that treatment with hydrochloric acid or sodium hydroxide caused a substantial reduction in the rise of the copolymer baseline, especially with sodium hydroxide, which also was accompanied by more intense cleavage product peaks.

In a second group of experiments the copolymer was heated without solvent and in solvent alone. The GPC and HPLC results for heating at 100°C without solvent, 4 h at 65°C in methanol, and 4 h at 100°C in water (Expts 5-7) are shown in Fig. 2.

**Fig. 2.**
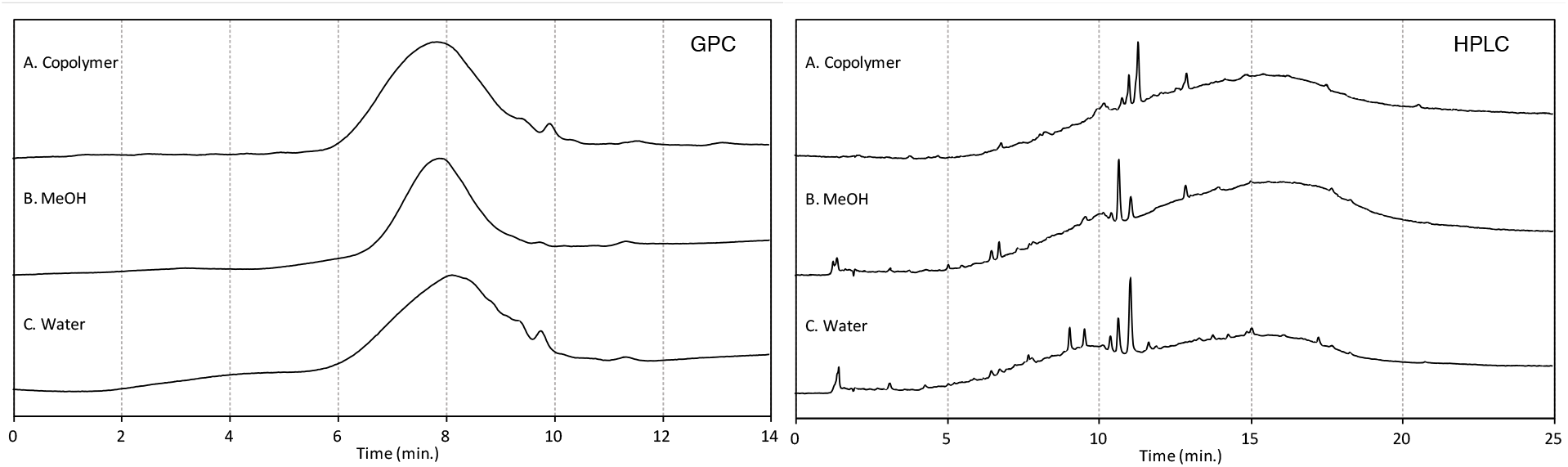
GPC-UV and HPLC-UV chromatograms at 220 nm of copolymer heated for 4 h at 100°C without solvent (A), 65°C in methanol (B) and 100°C in water suspension (C). (Expts. 5-7, Table 1). Injected sample sizes were ~200 μg.

Heating the copolymer at 100°C for 4 h without solvent caused only a small amount of decomposition, as can be seen by comparing Figs. 1A and 2A. However, in the presence of solvent there was some decomposition. Heating for 4 h in methanol (Fig. 2B) induced more decomposition than occurred during 14 d at room temperature (Fig. 1B), which was most evident in the comparison of the HPLC chromatograms. More extensive decomposition occurred in the water suspension heated at 100°C for 4 h. (Fig. 2C). The copolymer peak elution times were increased for treatment in methanol and more so for water, consistent with a greater degree of copolymer decomposition reducing mean polymer molecular weights.

In a third group of experiments, the copolymer was heated for 4 h at 65°C in methanol solutions of hydrochloric acid, sodium hydroxide and hydrogen peroxide, respectively (Expts. 8-10). The GPC and HPLC results are shown in Fig. 3.

**Fig. 3.**
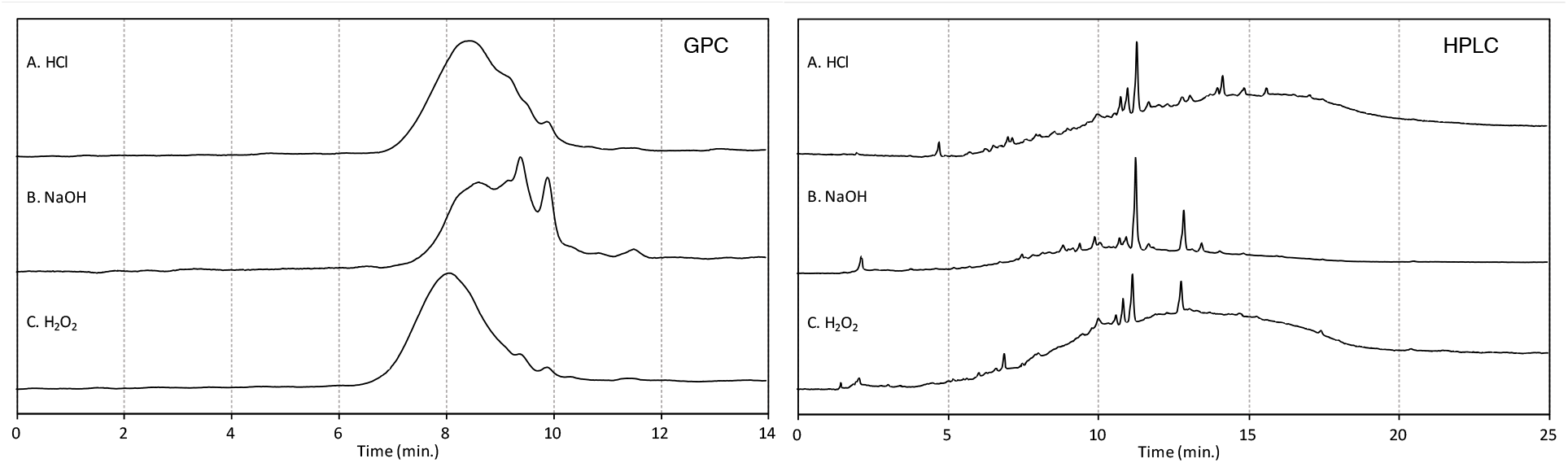
GPC-UV and HPLC-UV chromatograms at 220 nm of copolymer heated in methanol for 4 h at 65°C with excess hydrochloric acid (A), sodium hydroxide (B) and hydrogen peroxide (C). (Expts. 8-10, Table 1) Injected sample sizes were ~200 μg.

Comparison of the results for treatment with hydrochloric acid (Fig. 3A) vs. standing for 14 d at room temperature (Fig. 1D) revealed more extensive but still limited polymer breakdown. Treatment with sodium hydroxide showed increased, more extensive polymer breakdown (Fig. 3B vs. Fig. 1E), as shown by the overall change in the GPC profile and the significant reduction in the characteristic broad rise in the HPLC baseline (Fig. 3B).

Heating with hydrogen peroxide had little effect on the decomposition of the copolymer (Fig 3C).

Heating at 65°C with sodium hydroxide in methanol also was carried out with a larger excess of reagent or for a longer period of time (Expts. 11, 12). The GPC and HPLC results (not shown) indicated essentially no further change compared to treatment at the same temperature for 4 h or with less reagent (Fig. 3B).

Product recoveries for each experiment are given in Table 1. Treatment of the copolymer by standing for 14 d at room temperature in methanol (Expt. 1) or by heating, either without solvent (Expt 5) or in methanol (Expt 6), resulted in full recoveries. Recoveries were ~10% less for treatment in boiling water (Expt. 7) and even less after heating with sodium hydroxide or hydrochloric acid (Expts. 8, 9, 11, 12).

Copolymer dissolved in methanol containing hydrochloric acid had a deeper yellow colour than in methanol alone and became orange over time. A deep red colour developed in the treatment with sodium hydroxide in methanol. The aqueous phase obtained after ethyl acetate extraction of the sodium hydroxide-treated copolymer was orange, indicating the presence of water-soluble, ethyl acetate-insoluble breakdown products. The existence of water-soluble products not extracted by the workup process into ethyl acetate likely accounts for the reduced product recoveries.

### Breakdown Products

The identification of breakdown products made use of the fact that when introduced into the heated injector port of the GC-MS the OxBC copolymer decomposes thermally into several readily identified apocarotenoid breakdown products.^21^ This is illustrated by the GC-MS chromatogram in Fig. 4A. Eleven breakdown product structures were identified (structures **1-11**, Fig. 5).

**Fig. 4.**
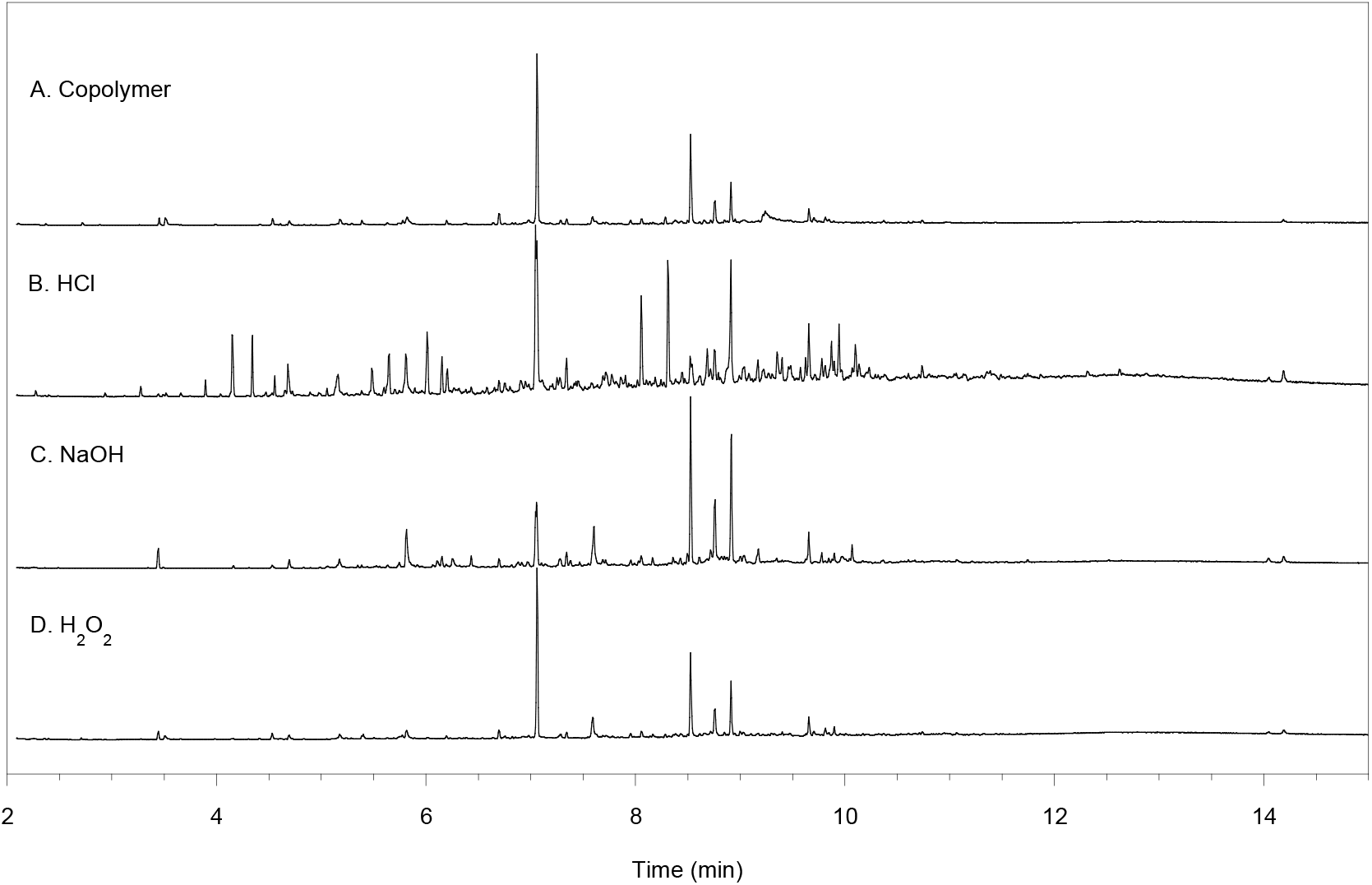
GC-MS chromatograms of unreacted copolymer (A) and of products after treatment in methanol for 4 h at 65°C with excess hydrochloric acid (B), sodium hydroxide (C) and hydrogen peroxide (D). (Expts. 8-10, Table 1.) The main peaks in A were identified previously.^21^

**Fig. 5.**
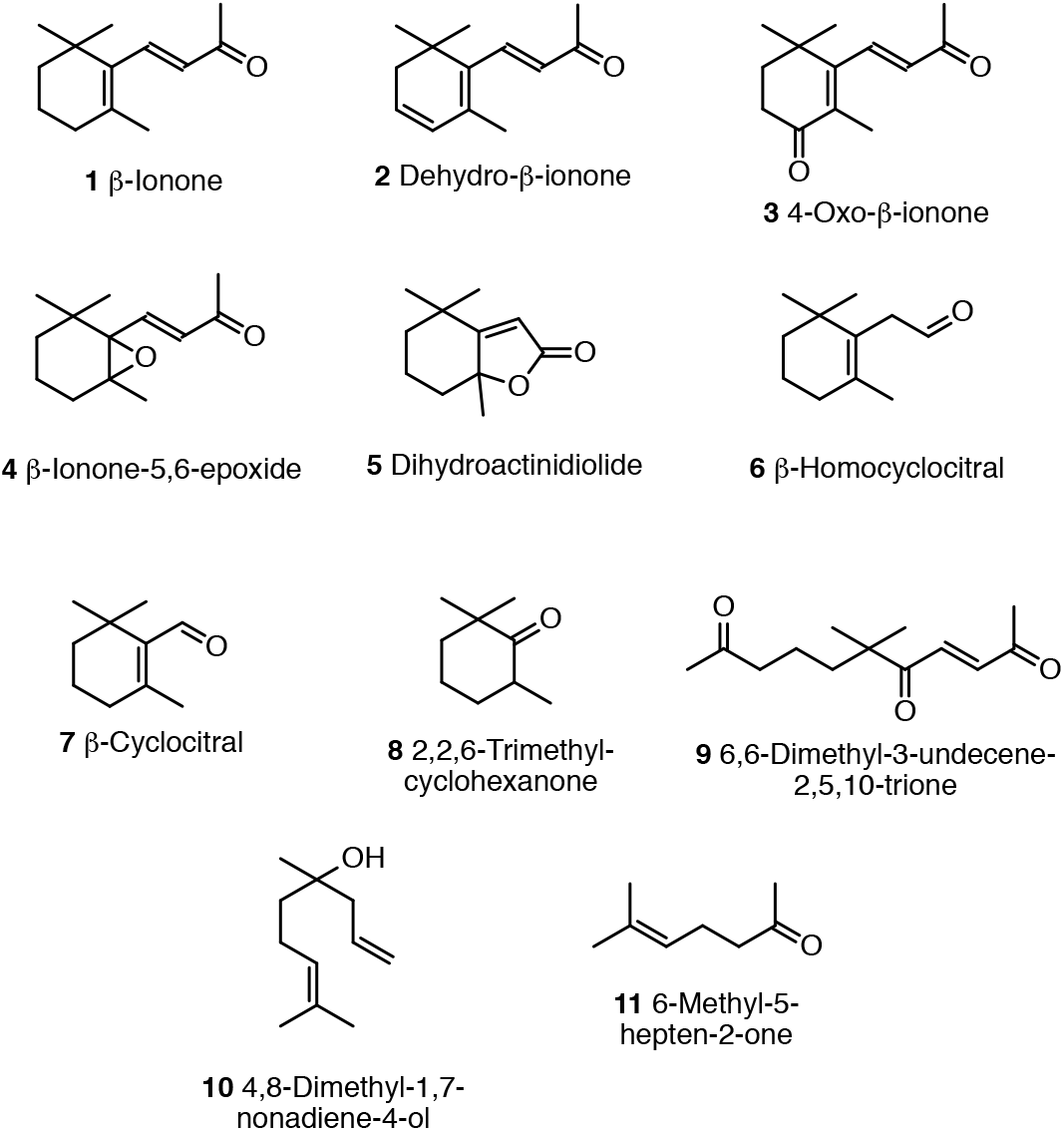
Copolymer breakdown structures **1-11** identified by GC-MS mass-spectral library matches after thermolysis of the copolymer compound during injection into the GC-MS injector port held at 250°C.

The presence of additional peaks in the GC-MS analysis of treated copolymer samples indicates the liberation of additional products from the copolymer.

The GC–MS chromatograms of the products obtained after standing for 14 d in methanol, with or without acetic acid, were similar to that for the copolymer by itself (results not shown). However, treatment with hydrochloric acid and sodium hydroxide, particularly when heated, gave more complex chromatograms, indicating additional breakdown products had formed (Figs. 4B, 4C).

Because of the possibility that some carboxylic acid breakdown products extracted into ethyl acetate may not be readily detected by GC-MS, extracted product mixtures from copolymer samples treated under acidic, basic or oxidizing conditions were subject to methylation with trimethyloxonium tetrafluoroborate. The methylated hydrochloric acid-treated copolymer was found to be especially rich in breakdown products (Fig. 6A). Many of these were esters, acetals and/or ketals that GC-MS analysis showed had already formed during treatment with methanolic hydrochloric acid.

**Fig. 6.**
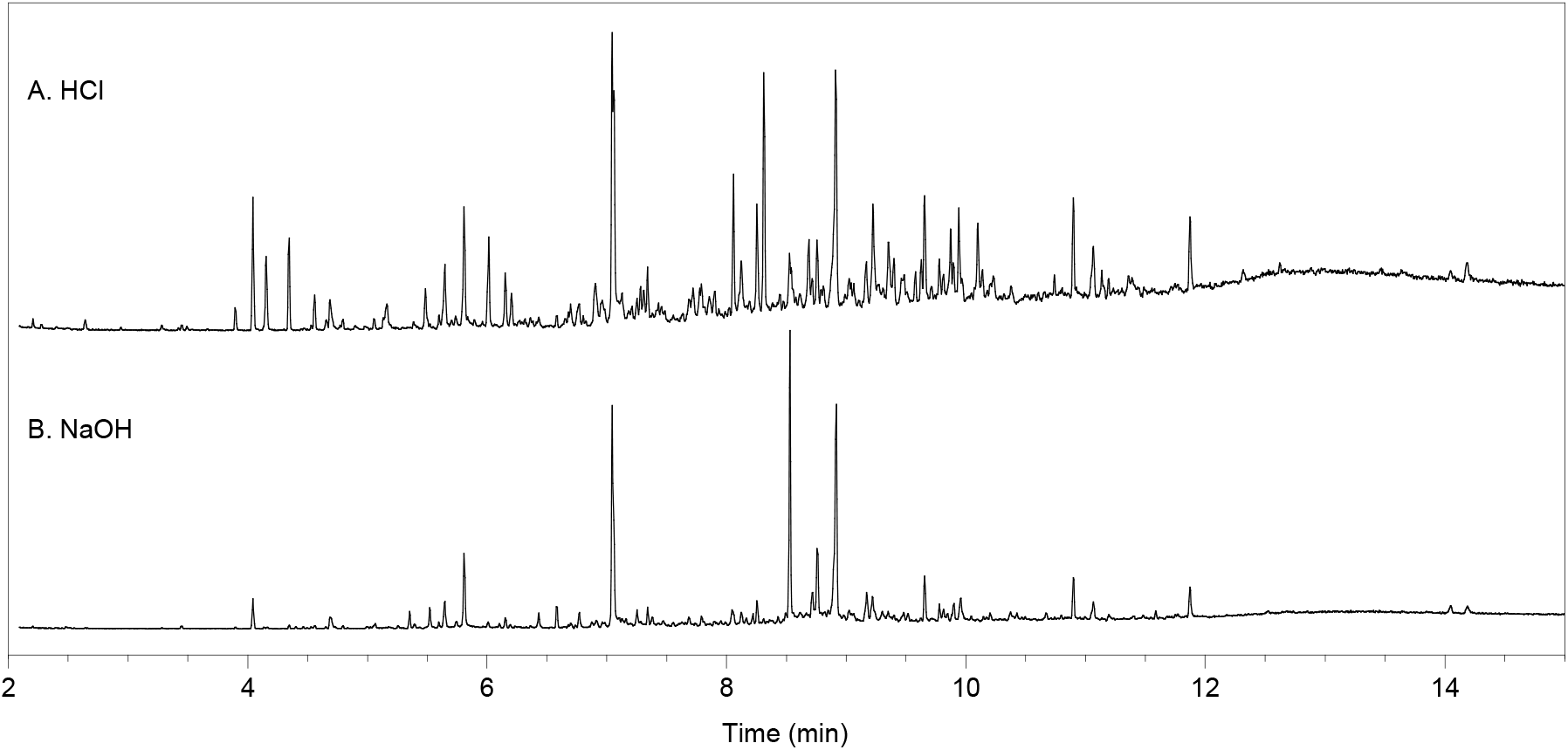
GC-MS chromatograms of copolymer products after treatment in methanol for 4 h at 65°C with excess hydrochloric acid (A) or sodium hydroxide (B) followed by methylation with trimethyloxonium tetrafluoroborate.

The GC–MS chromatograms of the product mixtures obtained from copolymer subjected to acidic and basic conditions were analyzed using GC-MS mass spectral library matching. Structures of an additional 34 compounds were identified with a mostly greater than 50% library match (structures **12-45**, Fig. 7). It was estimated that another 90 or more additional unidentified products were present, mostly at very low levels.

**Fig. 7.**
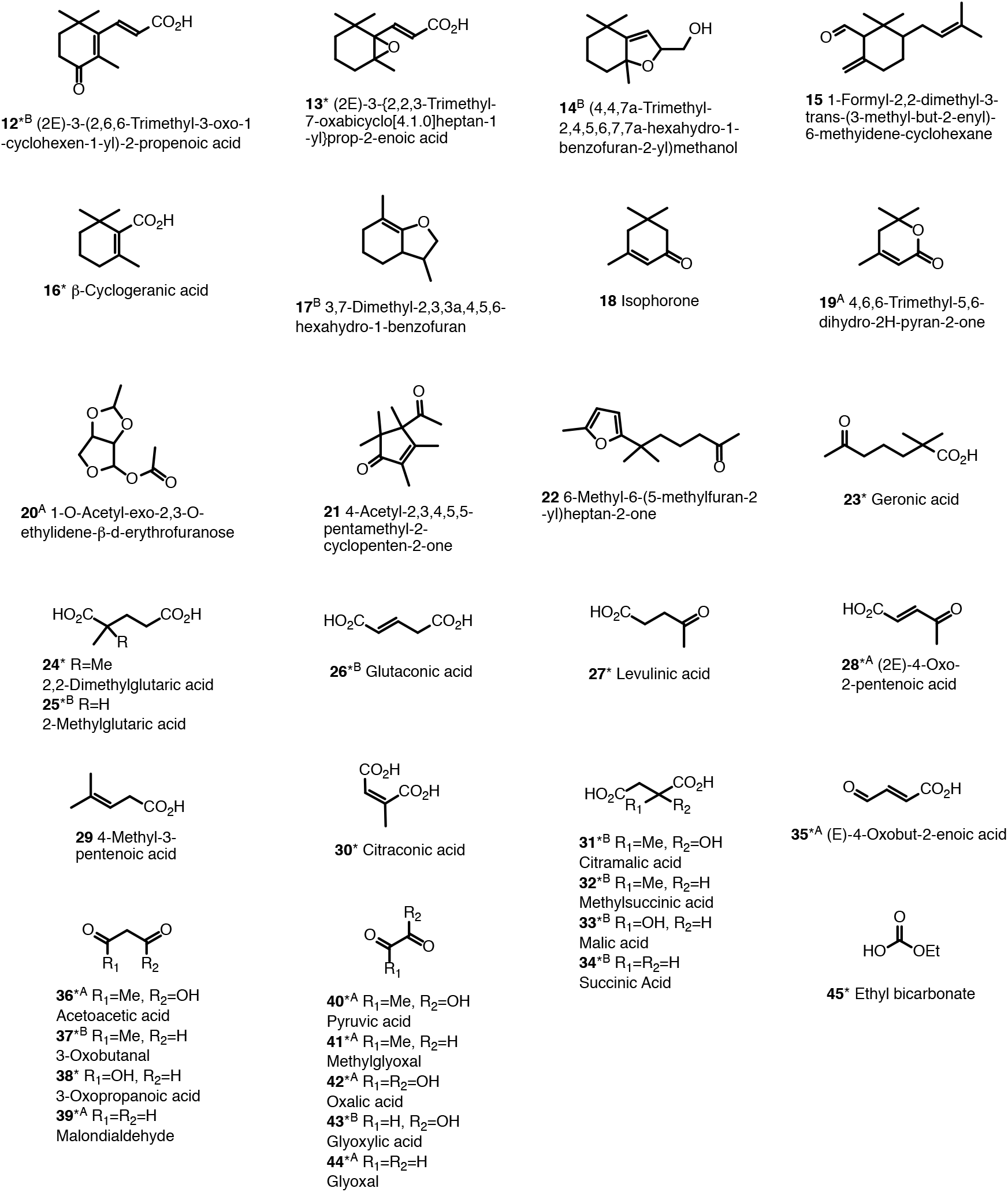
GC-MS mass-spectral library matches of 34 product structures, **12-45**, identified after hydrolysis of copolymer compound with hydrochloric acid or sodium hydroxide in methanol, in addition to the structures **1-11** identified in Fig. 5. Structure numbers bearing an asterisk are precursors of identified compounds, including methyl esters, acetals and/or ketals. Compounds obtained in common under both acidic and basic treatments are shown, unless indicated otherwise with a superscript A (acid) or B (base).

### Ozonolysis

Ozonolysis completely degraded the copolymer, as shown by the absence of both its characteristic peak in the GPC and the distinctive large baseline rise in the HPLC (results not shown). Also, the GC-MS chromatogram of the ozonolysis product was distinctly different from that of the pure copolymer (results not shown).

The colourless character of the product was consistent with the absence of residual conjugated double bonds, replaced very likely with carboxylic acid and keto groups. Indeed, GC-MS analysis of both the reaction product and of a sample esterified with trimethyloxonium tetrafluoroborate confirmed the presence of carboxylic acids. In addition to structures **23-25, 28-30** and **33** (Fig. 7), structures **46-58** were identified by mass-spectral library matching (Fig. 8).

**Fig. 8.**
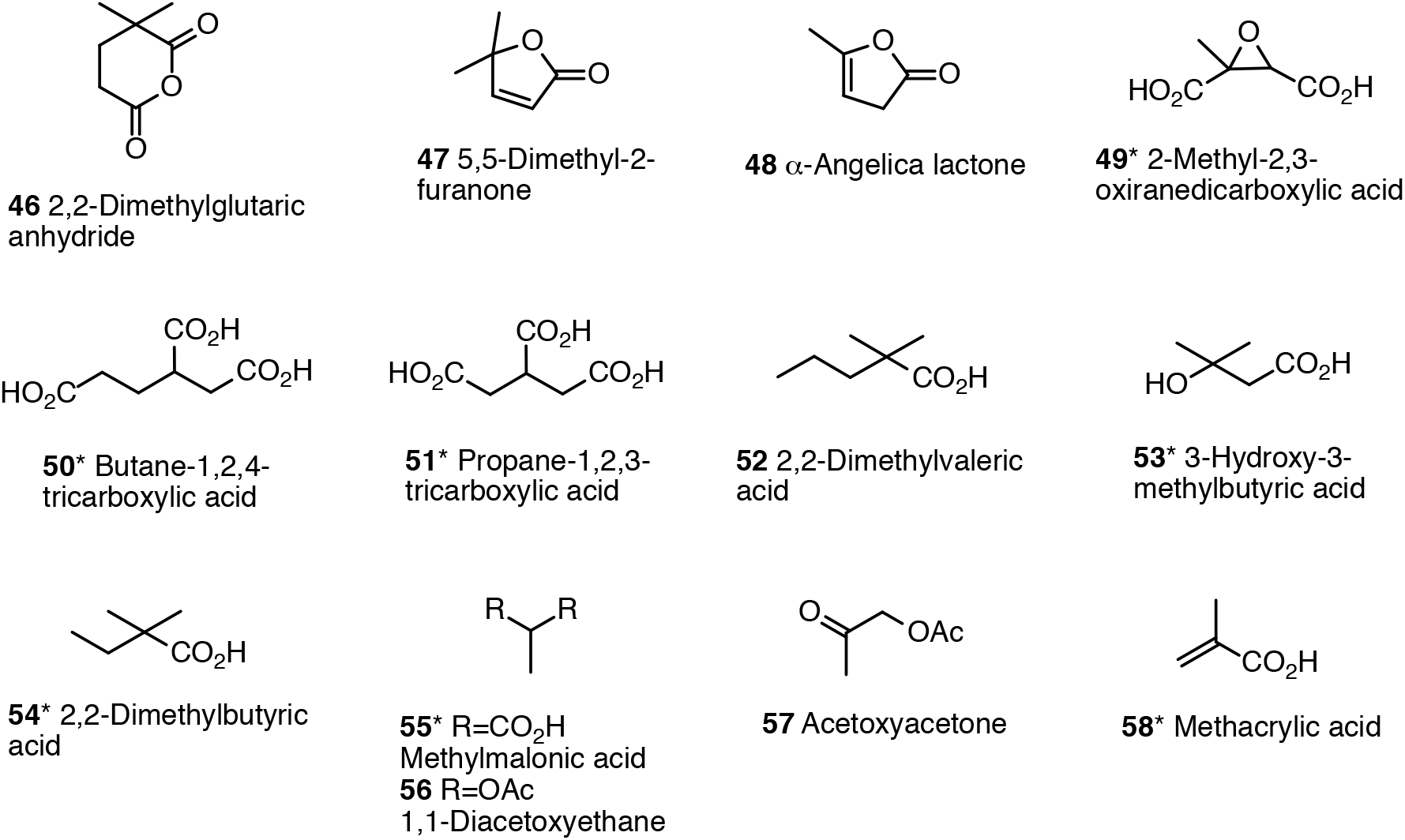
Structures **46-58**, in addition to structures **23-25, 28-30** and **33**, identified by GC-MS mass-spectral library matches of products recovered after ozonolysis of the copolymer compound. Structure numbers bearing an asterisk are precursors of identified methylated compounds obtained by reaction with trimethyloxonium tetrafluoroborate.

Because of some chemical similarities with the copolymer noted earlier,^5, 21^ a sample of sporopollenin from lycopodium was ozonized and analyzed in a similar manner. Product structures identified by GC-MS mass-spectral library matches after methylation with trimethyloxonium tetrafluoroborate are shown in Fig. 9.

**Fig. 9.**
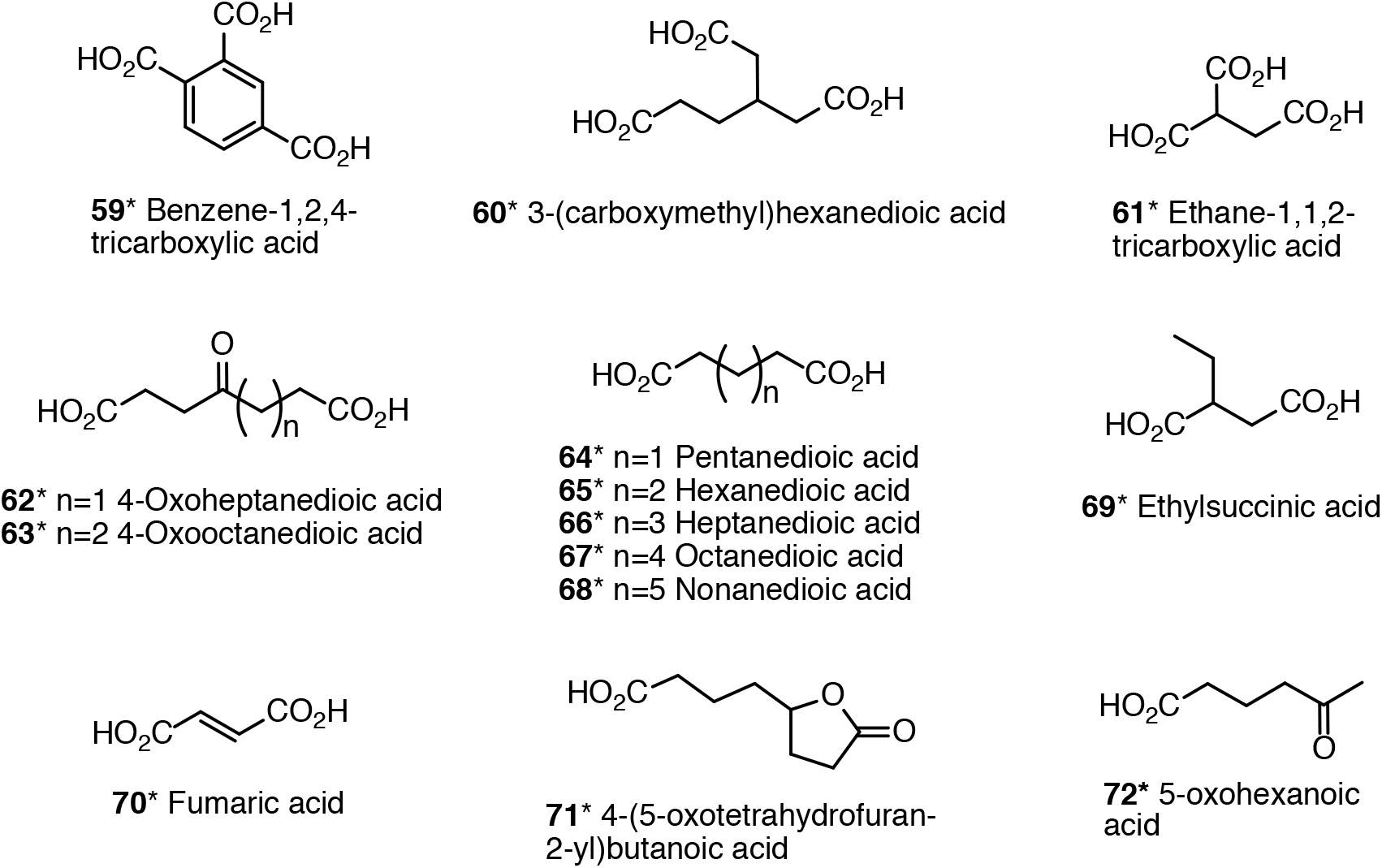
Structures **59-72**, in addition to structures **25**, **26**, **28**, **31-34** (Fig 7), **50**, **51** and **55** (Fig. 8), identified by GC-MS mass-spectral library matches of products recovered after ozonolysis of lycopodium sporopollenin, followed by methylation with trimethyloxonium tetrafluoroborate. Structure numbers bearing an asterisk are precursors of identified methylated compounds.

### GA content of copolymer breakdown product mixtures

The results of the GA analyses are given in Table 2 where the quantity of GA is expressed relative to the starting quantity of copolymer (%, w/w). It is apparent that GA exists in the copolymer in more than one form and GA release occurs with varying degrees of ease from the individual forms.

**Table 2.** Geronic acid content of OxBC copolymer breakdown products after various treatments.

| Treatment | Experiment | Geronic Acid<sup>a</sup>, % |
| --- | --- | --- |
| None | - | 0.29 |
| Heat, 100°C, 4 h (no solvent) | 5 | 0.32 |
| Reflux, ethyl acetate, 77°C, 4 h | - | 0.32 |
| Reflux, methanol, 65°C, 4 h | 6 | 0.32 |
| Reflux, H<sub>2</sub>O, 4 h | 7 | 0.44 |
| Reflux, methanol with HCl (0.12 M), 4 h | 8 | 0.42 |
| Reflux, methanol with NaOH (0.12 M), 4 h | 9 | 0.57 |
| Reflux, methanol with aq. H<sub>2</sub>O<sub>2</sub> (0.12 M), 4 h | 10 | 0.48 |
| Reflux, methanol with NaOH (0.12 M), 24 h | 11 | 0.58 |
| Ozonolysis in acetic acid, 6 h, aq. H<sub>2</sub>O<sub>2</sub> work up | 13 | 1.42 |

<sup>a</sup> Relative to initial copolymer (w/w).

The unreacted copolymer yielded 0.29% GA, indicating that either during the analytical methylation reaction GA was released from the copolymer or GA had not been completely removed during copolymer isolation by hexanes precipitation from ethyl acetate solution. GA could be non-covalently bound by hydrogen bonding via the copolymer’s rich endowment of polar oxygen functionalities.

Refluxing the copolymer in ethyl acetate, methanol or heating in isolation to 100°C for 4 h yielded only a slight increase in GA, to 0.32%. However, the GA yield increased to 0.44% when a suspension of the copolymer was heated for 4 h in boiling water.

The GA yield was 0.42% for reflux with methanolic hydrochloric acid and was almost doubled to 0.57% for reflux with methanolic sodium hydroxide. There was little difference between heating with sodium hydroxide for 4 h, 24 h or for 4 h with 4-fold the amount of sodium hydroxide, indicating a limit to the GA produced under basic conditions.

Refluxing with aqueous hydrogen peroxide in methanol increased the GA yield to 0.48%, which may have been due to water itself and/or the oxidizing effect of hydrogen peroxide.

Ozonolysis of the polymer gave the largest yield of GA (1.42% total), confirming the presence of moieties within the copolymer that are converted to GA upon further oxidation. When combined, these sources yield an extra 1.1% GA (1.42%, ozone treatment - 0.29%, isolated copolymer).

## Discussion

Copolymerization with oxygen lies at the heart of understanding the course and outcome of β-carotene autoxidation. The emerging picture is that the apocarotenoid cleavage products are generated by the polymerization process itself and possibly, subsequently, from limited decomposition of the resulting copolymer product.

As Scheme 1 illustrates, spontaneous free radical oxidation starts with an O_2_ molecule adding to the β-carotene molecule. The intermediate oxygen adduct radical can either cleave into two apocarotenoid products or, preferentially, undergo further addition of an O_2_ molecule at various sites of the remaining conjugated double bonds over which the unpaired electron is delocalized. The high degree of polyene unsaturation remaining in the early oxygen radical adducts strongly favors competitive addition of O_2_ over cleavage. Progression through successive intermediates leads to the copolymer as the major product and, in lesser amount, the apocarotenoids. The present study shows the resulting copolymer compound is complex, being made up of precursor chemical constituents that are capable of giving rise to a diverse range of many small apocarotenoids (< C_20_).

In Scheme 1 the apocarotenoid products formed during the oxidation arise from scission reactions of the copolymer radical rather than by the conventionally understood direct oxidative cleavage of individual double bonds in the polyene backbone.^5^ The larger apocarotenoids that are formed early in the reaction, by nature of their remaining high unsaturation, are still highly reactive towards the growing copolymer radical intermediates, or even oxygen itself, and re-enter the chain reaction, undergoing further oxidative conversions. The products that subsequently emerge as the polymerization process advances are the smaller apocarotenoids, most of which have undergone further oxidative and other secondary chemical transformations.

This study shows that the copolymer itself can be a source of many apocarotenoid and secondary breakdown products under certain conditions. It remains to be seen to what extent copolymer breakdown occurs during metabolism in plants and animals.

### Copolymer stability and breakdown products

Heating the copolymer alone at 100°C and heating with strong acid or base in methanol tested the stability of the copolymer and its ability to release its constituent chemical entities.

GPC and HPLC analyses have shown the copolymer, without solvent, is resistant to decomposition to at least 100°C for a minimum of 4 h (Fig. 2A) but does quickly decompose into apocarotenoids at a temperature of ~250°C in the GC-MS injector port (Figs. 4A, 5). The compound is susceptible to partial chemical breakdown by hot water and more so by strong acid and base. Heating with sodium hydroxide produced the largest change. Substantial amounts of a lower molecular weight polymeric material were still present, albeit in some apparently altered form that did not release further breakdown products.

Treatment with hydrogen peroxide tested the possibility of an oxidative degradation pathway, as could conceivably occur *in vivo* through interaction with white blood cells, for example. However, the reagent had little effect.

The incomplete recoveries of products extracted into ethyl acetate from the sodium hydroxide and hydrochloric acid copolymer treatments (Table 1), especially after heating, together with the appearance of color in the aqueous fraction during workup, indicated the generation of new, water-soluble, ethyl acetate-insoluble products. The nature of these products has not been examined.

GC-MS analyses of the chemical breakdown products showed the copolymer is a latent source of many small apocarotenoids and secondary products. Forty-five structures were identified. A few structures (e.g., **20, 21**, Fig 7) cannot be entirely certain using mass spectral matching alone. Another 90 or more mostly minor GC-MS peaks had unassigned structures. These peaks may have included several structures reported in an earlier detailed chemical analysis of the individual apocarotenoids in the OxBC low molecular weight fraction that were not identified in the copolymer breakdown products.^5^ These compounds included 2-methyl-6-oxo-2,4-heptadienal, 2-hydroxy-2,6,6-trimethylcyclohexanone, 4-oxo-β-cyclocitral, β-ionylidene acetaldehyde, β-ionylidene acetaldehyde-5,6-epoxide, 3-(2,2,6-trimethyl-7-oxabicyclo[4.1.0]heptan-1-yl)oxiran-2-yl acetate, and β-apo-13-carotenone-5,6-epoxide.

Various carotenoid-derived apocarotenoids are already known for their flavour and fragrance properties.^6^ They are found, for example, in numerous leaf products, such as tea and tobacco, essential oils, fruits, vegetables, spices, wine, rum, coffee, honey and seaweed.^27^ Thirteen of the identified copolymer-derived structures, **1, 4-6, 7, 8, 11** (Fig. 5), **18, 27, 33, 34, 40** and **41** (Fig. 7) are currently listed as Generally Recognized as Safe (GRAS) flavor agents.^7^

Eight of the identified volatile apocarotenoids, in particular compounds **1, 2, 3, 4, 5, 7, 11** and **16**, have been detected in plants by head space analysis of Arabidopsis leaves.^28^

The breakdown products liberated by acid or base treatment include numerous small molecule carboxylic acids (Fig. 7: **23-36, 40, 42, 43**), several of which appear not to have been known as β-carotene oxidation products. These include keto carboxylic acids (**23, 27, 28, 35, 36, 38** and **40**) and dicarboxylic acids (**24-26, 30-34** and **42**), several of which are known central metabolic intermediates (e.g., pyruvic, glyoxylic and succinic acids). These and other secondary compounds that are not simple cleavage compounds have undergone further chemistry that is typical of peroxide rearrangements, e.g., the Baeyer-Villiger rearrangement.^29^

Of note are the short dialdehyde compounds, methyl glyoxal (**41**) and glyoxal (**44**). A recent model study of the rates of biosynthesis and degradation of carotenoids in plant tissue has found that oxidative β-carotene degradation occurs mostly non-enzymatically and methyl glyoxal and glyoxal are putative end-product metabolites.^15^ Increased levels of β-carotene correlated with increased levels of these compounds and, furthermore, increased levels of geronic acid, a marker compound for the β-carotene copolymer.^5^ The present study supports the copolymer or its intermediates as potential sources of these two dialdehyde metabolites.

The formation of methyl glyoxal takes on additional significance with the recent report that Manuka honey-derived methylglyoxal can enhance microbial sensing by mucosal-associated invariant T cells, which are found, for example, in the oral and gastrointestinal tracts.^30^ Methyl glyoxal can activate these cells, leading to the production of a variety of cytokines, chemokines and responses that can in turn protect the body against microbial infection and potentially support immune homeostasis.

### Ozonolysis

The complete oxidative breakdown by ozonolysis of the copolymer and GC-MS mass-spectral matching of the products further confirmed the extraordinary complexity of its chemical makeup (Fig. 8). The ozonolysis of sporopollenin was carried out to compare product profiles (see Fig. 9). Previously, we had suggested, on the basis of the high oxygen content and FTIR data, a chemical similarity existed between the copolymer and sporopollenin.^5, 21^ Comparison of structures identified in Figs. 7, 8 and 9 show that although there are several structures in common (**25, 28, 33, 50, 51** and **55**), there are many more that do not share structures in common. In particular, it is difficult to imagine how benzene 1,2,4-tricarboxylic acid, **59**, and the unbranched alkyl dicarboxylic acids **62-68** from sporopollenin could be formed from the polyisoprenoid structure of β-carotene. Furthermore, the first ever detailed structure of a sporopollenin, published recently,^31^ revealed there is no carotenoid or isoprenoid component in sporopollenin’s complex structure.

### The copolymer as a source of GA

GA is one of the most abundant apocarotenoids in OxBC.^5^ The results of the GA analyses of the copolymer breakdown products provide further support for the conclusion that OxBC apocarotenoids and secondary products originate from the copolymer and/or its intervening intermediates. Table 2 shows the copolymer is a source of GA beyond the free GA already present in the apocarotenoid fraction of OxBC. A small amount of GA is readily available directly from the copolymer, without treatment, and does not increase under mild conditions. Under more vigorous conditions, the amount is increased by reflux in water, methanolic hydrochloric acid and methanolic sodium hydroxide. With ozone, a larger, roughly 4-fold increase occurs, from 0.3% to 1.4%. Previously, an extended oxidation of β-carotene with pure oxygen at room temperature over 7 days also increased the amount of GA over the usual amount generated over the 1 day period used to prepare OxBC in pure oxygen.^5^

In a preliminary, ongoing oxidation study of β-carotene carried out with 99% pure ^18^O_2_, it was discovered that the free GA in the low molecular weight OxBC apocarotenoid fraction contains ^18^O in only two of the three GA oxygen atoms. Both carboxyl oxygens contain ^18^O, whereas the keto oxygen contains ^16^O. This result is interpreted to mean that GA is produced from an ^18^O-labeled precursor in the copolymer intermediates during reaction by hydrolysis with H_2_^16^O present in the ethyl acetate solvent. The study will be reported elsewhere when complete.

### Potential significance of the copolymer

In plants, a function has emerged for β-carotene as a source of β-apocarotenoid products that are generated in response to environmental oxidative stresses.^11, 17^ When plants are exposed to light in excess of their photosynthetic capacities, reactive oxygen species (ROS), especially singlet oxygen (^1^O_2_), can cause lipid peroxidation that generates toxic reactive species. Under such circumstances, β-carotene can act as a chemical quencher of ^1^O_2_. The generation of very small quantities of a variety of β-apocarotenoid products, some of which are reactive electrophilic species, can induce gene expression changes that ultimately result in tolerance to oxidative stress conditions. β-Carotene oxidation therefore represents an early warning signal of oxidative light stress in plants. β-Cyclocitral (**7**) and dihydroactinidiolide (**5**), are recognized signaling compounds that at very low concentrations (ng/g) activate defenses against ROS.^12, 17^ Cyclogeranic acid (**16**) can induce protection against the effects of drought.^17^

In non-photosynthetic plant tissues, Schaub and co-workers^15^ and Koschmieder and coworkers^16^ have utilized a model system of transgenic Arabidopsis roots over-accumulating β-carotene and β-apocarotenoids to demonstrate there is rapid turnover of β-carotene that principally involves non-enzymatic oxidation of β-carotene. In recognizing that many apocarotenoids are reactive electrophiles with α,β-unsaturated carbonyl moieties, it was shown by transcriptome analyses of the Arabidopsis roots that apocarotenoids are metabolized by enzymes known for detoxification of xenobiotics and reactive carbonyl species.^16^ Schaub et al., have noted that in stored plant tissues the amount of degraded β-carotene exceeded the concentrations of apocarotenoids present by several orders of magnitude.^20^ The copolymer product was identified as being a major portion of the degraded β-carotene.

Given that β-carotene autoxidation proceeds by free radical copolymerization with oxygen regardless of environment, it is highly likely that in aerobic plant cells oxidation of β-carotene also will generate copolymers, regardless of what reactive species initiates the oxidation. This provides a means to safely and substantially reduce the levels of potentially toxic *free* apocarotenoids by sequestration in chemically bound forms in the relatively stable copolymer form. The levels of free apocarotenoids are therefore kept at the low levels required to be able to act as signals to maintain cellular defences against ROS. The copolymer stability results presented here do not preclude the possibility that under oxidative stress conditions the polymer could be partially metabolized to release further apocarotenoids as the need arises.

In animals a similar role for the copolymer in controlling and limiting the concentration of free, reactive apocarotenoids can be plausibly envisaged, assuming non-enzymatic autoxidation pathways for β-carotene degradation exist in animals. By analogy to its role in plants, β-carotene could serve as an early warning system for potentially toxic ROS levels in animal cells. Although present in low concentration in cells, the highly reactive β-carotene still is much more susceptible to reaction with ROS than other, more abundant, less reactive membrane lipids and therefore would be oner of the first lipid sites of attack. The small apocarotenoid products generated could play a role similar to that in plant cells, by inducing similar mechanisms of ROS protection. Model transcriptome studies using OxBC, in a manner somewhat similar to those conducted with Arabidopsis plants, could help uncover any potential involvement of apocarotenoids as signals of ROS.

The copolymer itself has biological activity in mammalian cells. In human THP-1 monocyte cells the membrane content of CD-14 immune surveillance receptors was significantly increased to the same level as that of the parent OxBC, whereas the low molecular weight fraction containing the apocarotenoids showed little activity.^25^

The OxBC product provides a ready source of both copolymer and apocarotenoids that, in principle, are available for distribution to tissues, thereby circumventing or supplementing the need for products from prior oxidation of β-carotene *in situ*. A study in mice orally administered OxBC is presently underway to determine if the OxBC copolymer is transported from the gut into the circulation. This study is in support of understanding how OxBC provides systemic health benefits as, for example, in a dairy calf model of bovine respiratory disease-induced lung inflammation^26^ and reduction of mastitis in dairy cows.^24^ It is likely that the free apocarotenoids in orally administered OxBC would be rapidly removed from the circulation by metabolism, e.g., conjugation,^16^ which reduces the possibility they would be sufficiently available for sustained effect in more remote tissues. Is the copolymer therefore more persistent in the circulation and therefore more available for uptake into tissues and thereby acting as a vehicle for transporting apocarotenoids for subsequent release into tissues? Release may be more facile at sites of inflammation, given the various enzymatic and oxidative activities associated with that condition. Or does the copolymer itself exert a dampening effect upon inflammation?

## Acknowledgments

Dr Julian Koschmieder is thanked for helpful discussions.

---

**Scheme 1.**
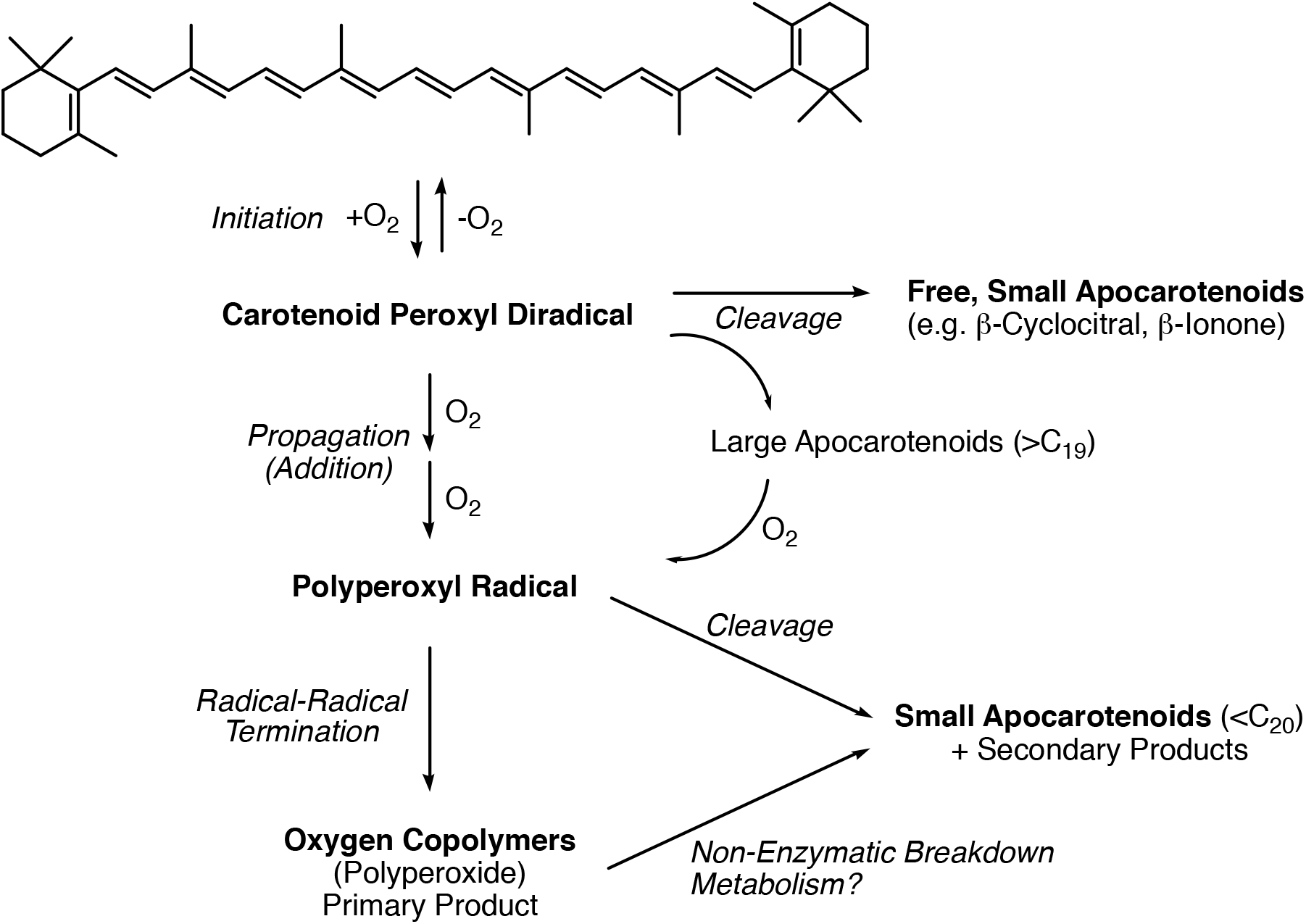
Mechanism of β-carotene oxidation. O_2_, itself a diradical, reversibly adds to β-carotene to form a peroxyl diradical intermediate that then enters a chain reaction by adding further O_2_, or it may undergo cleavage at this or later intermediate stages to release apocarotenoid molecules. The larger, still highly unsaturated apocarotenoid cleavage products that are initially formed can eventually re-enter the chain reaction as the reaction progresses. The growing oxygen copolymer radical also undergoes scission reactions to give small apocarotenoid and secondary products, ultimately terminating in a relatively stable copolymer product with a molecular weight ranging from several hundred to approximately 8,000 Da.

